# IL-17 sensitises sensory neurons and colonic afferents to noxious stimuli in a PI3K dependent manner

**DOI:** 10.1101/2025.04.16.649143

**Authors:** Luke W Paine, James P Higham, Katie H Barker, Sofia Pavlou, Fraser Welsh, Ewan St. John Smith, David C Bulmer

## Abstract

Managing visceral pain associated with gastrointestinal (GI) disease remains a significant challenge due to the gut-related side effects and contraindicated use of many commonly used painkillers in people with inflammatory bowel disease (IBD). Consequently, it is crucial to deepen our understanding of the mediators and mechanisms underlying inflammatory pain in people with IBD. To do this, we compared bulk RNA sequencing data from colonic biopsy samples from people with IBD with single-cell RNA sequencing data from colon projecting dorsal root ganglion (DRG) neurons in mice to generate an interactome of putative pro-nociceptive cytokine signalling pathways. This *in silico* analysis revealed a 10-fold increase in *IL17A* expression in samples from people with ulcerative colitis (UC) alongside marked co-expression of *Il17ra* with *Trpv1* in colon-projecting DRG neurons in mice, highlighting a likely role for interleukin-17 (IL-17) in colonic nociceptor signalling in people with UC. In support of this, Ca^2+^ imaging studies demonstrated that IL-17 stimulates DRG sensory neurons co-sensitive to capsaicin with a similar proportion responding in neuron-enriched cultures generated by magnetic-activated cell sorting, thus confirming that IL-17 directly activates DRG neurons.

IL-17-evoked Ca^2+^ signals were attenuated by TRPV1 inhibition, consistent with nociceptor activation, and blocked by inhibition of phosphoinositide 3-kinase (PI3K) activity, consistent with the known role for PI3K as a downstream effector of IL-17 receptor signalling. In keeping with these observations, IL-17 enhanced murine colonic afferent responses to colorectal distension at noxious distension pressures, an effect also blocked by PI3K inhibition. Overall, these findings demonstrate a pro-nociceptive effect of IL-17 in the GI tract, thus highlighting the potential utility of IL-17-targeting therapies to reduce pain in people with UC.

## Introduction

Persistent inflammation in the gastrointestinal (GI) tract gives rise to a range of colitides, collectively referred to as inflammatory bowel disease (IBD) (C. Abraham & J. H. Cho, 2009). A hallmark of these diseases is the elevated production of proinflammatory cytokines which, alongside driving inflammation, are increasingly recognised as playing a key role in the development of visceral hypersensitivity and chronic abdominal pain due to their direct effects on nociceptor signalling (Friedrich et al., 2019; Grundy et al., 2019).

Selectively sequestering IL-23 has proven successful in reducing inflammation associated with psoriasis (Hawkes et al., 2018) and offers an alternative to the widely prescribed anti-tumour necrosis factor (TNF) sequestering therapies commonly used in the treatment of IBD (Gaffen et al., 2014). IL-23, a member of the IL-12 cytokine family, is released by infiltrating macrophages and monocytes recruited to inflamed sites and plays a critical role in the development of chronic inflammation due to its ability to drive IL-17 production from T-helper type 17 (Th17) cells (Langrish et al., 2005; McGeachy et al., 2009) and innate lymphoid cells (Buonocore et al., 2010).

The IL-17 cytokine family comprises six subtypes (IL-17A-F), each with distinct expression patterns and signalling properties. IL-17A is the most well-characterised subtype, typically forming covalent homodimers of 31 kDa, although heterodimerisation with other IL-17 subtypes has been reported (Chang & Dong, 2007). IL-17A exerts its effects through the widely expressed IL-17 receptor, a heterodimeric complex of IL-17RA and IL-17RC subunits (Toy et al., 2006). Functionally, IL-17A signalling triggers pro-inflammatory gene expression, akin to IL-1R activation, including the induction of nuclear factor-κB (NF-κB) and other transcription factors implicated in intestinal inflammation (Chang et al., 2006; Linden, 2007). Additionally, IL-17 regulates intestinal epithelial permeability (Lee et al., 2015) and amplifies TNF and IL-6 release (Gaffen et al., 2014), thereby positioning dysregulated IL-17 signalling a key contributor to IBD pathophysiology.

While the role of IL-17 in inflammation is well established, its contribution to the abdominal pain associated with GI diseases remains poorly understood. In models of arthritis, IL-17 has been shown to mediate mechanical pain, with its neutralization preventing methylated bovine serum albumin (mBSA)-induced articular hypersensitivity (Pinto et al., 2010) and intra-articular injection of IL-17 sensitising nociceptive C-fibres and enhancing the excitability of rat dorsal root ganglia (DRG) neurons (Richter et al., 2012).

More recently, behavioural, genetic, pharmacological and electrophysiological studies have further highlighted the importance of an IL-23/IL-17/TRPV1 signalling axis with the pro-nociceptive effects of IL-17 being abolished following TRPV1 blockade (Luo et al., 2021). While these findings stem from somatic pain models, it remains unclear whether similar mechanisms operate in the GI tract. As such, the goal of the present study was to confirm the stimulatory effect of IL-17 on capsaicin-sensitive sensory neurons and confirm the translation of these findings to colonic afferents sensitive to noxious colorectal distension.

## Materials and Methods

### Animals and Ethical Approval

Adult male C57BL/6J mice (8–16wk) were obtained from Charles River (Cambridge, UK; RRID: IMSR_JAX:000664). Mice were conventionally housed in temperature-controlled rooms (21°C) following a 12-hour light/dark cycle and provided with nesting material, a plastic shelter and unrestricted access to food and water. All animal experiments were conducted in compliance with Schedule 1 of the Animals (Scientific Procedures) Act 1986 Amendment Regulations 2012.

### Reagents

Stock concentrations of IL-17 (IL-17A, 7956-ML-025; H_2_O with 0.2% (w/v) bovine serum albumin), and capsaicin (1 mM; 100% ethanol) were obtained from Sigma-Aldrich and dissolved as instructed. SB203580 (10 mM; DMSO), A425619, AM0902 and thapsigargin (all at 1 mM; DMSO) were obtained from Tocris Bioscience and made up as described. Wortmannin (1 mM; DMSO) was purchased from Cayman chemicals. Atropine and nifedipine (100 mM; 100% ethanol and DMSO, respectively) were purchased from Sigma-Aldrich and dissolved as described. Immediately preceding the experiment, all drugs were diluted to their working concentrations in either extracellular solution (ECS) or Krebs buffer as appropriate.

### *In silico* analysis of bulk RNAseq of colonic biopsies and scRNAseq of murine colonic sensory neurons

For bulk RNAseq of colonic biopsies, data were analysed from (Higham et al., 2024). In brief, biopsies were collected from patients undergoing diagnostic colonoscopy at the Royal London Hospital. Biopsies were taken from inflamed sites of ulcerative colitis (UC) patients (n = 9 biopsies from N = 9 patients) and Crohn’s disease (CD) patients, the latter of which was divided into treatment-naïve (CDN; n = 8 biopsies from N = 7 patients) and treatment-refractory (CDT; n = 11 biopsies from N = 7 patients) subgroups. All patients had reported abdominal pain within the four weeks prior to endoscopy.

Biopsies were also obtained from the sigmoid colon of patients experiencing abdominal pain but without visible inflammation, later diagnosed with recurrent abdominal pain (RAP) (n = 21 biopsies from N = 16 patients). Additionally, samples were collected from non-inflamed (NI) individuals (n = 14 biopsies from N = 8 patients) who had neither reported abdominal pain nor exhibited inflammation upon endoscopy.

For single-cell RNA sequencing (scRNA-seq) of murine sensory neurons, data were obtained from (Hockley et al., 2019).

### Primary culture of mouse dorsal root ganglion neurons

Mice were euthanised using a rising concentration of CO_2_ followed by exsanguination and thoracolumbar DRG were collected from T12-L5 spinal segments, specifically chosen as they are the predominate DRG that innervate the distal colon (Robinson et al., 2004). The collected DRG were placed in ice-cold Leibovitz L-15 medium (containing GlutaMAX supplement and 2.6% (v/v) NaHCO_3_). DRG were then incubated at 37°C with type 1A collagenase (1 mg/ml) and trypsin solution (1 mg/ml) both with 6 mg/ml BSA for 15 and 30 minutes, respectively. Following this enzymatic digestion, DRG were resuspended in 2 ml dissociation media (containing Leibovitz L-15 medium, GlutaMAX supplement, 2.6% (v/v) NaHCO_3_, 10% (v/v) fetal bovine serum (FBS), 1.5% (v/v) glucose and 300 units/ml penicillin / 0.3 mg/ml streptomycin (P/S)) and mechanically triturated with increasing intensity to dissociate the DRG. The solution was centrifuged (100 *g*) and supernatant collected for five consecutive trituration’s. Following centrifugation (5 min, 100 *g*) and resuspension, the supernatant was plated (50 µl per dish) onto laminin coated (Thermo Fisher: 23017015) 35 mm poly-D-lysine-coated glass bottom dishes (MatTek P35GC-1.5-14-C) and incubated at 37°C for 2-3 hours to facilitate cell adhesion. Subsequently, the dishes were flooded with dissociation media and incubated overnight (37°C, 5% CO_2_).

### Ca^2+^ imaging

Extracellular solution (ECS) was prepared with the following composition (in mM): 140 NaCl, 4 KCl, 1 MgCl_2_, 2 CaCl_2_, 4 glucose, 10 HEPES. The pH was adjusted to pH 7.4 using NaOH and the osmolarity to 290-310 mOsm with sucrose. Cultured DRG sensory neurons were incubated with 100 µl Fluo-4-AM (10 µM in ECS, Invitrogen), a cell-permeable fluorescent Ca^2+^ indicator, for 30 minutes at room temperature, shielded from light. For inhibitor studies, cells were pre-incubated for 10 minutes with 200 µl of drug prior to imaging. For studies with Ca^2+^ omitted from the ECS (in mM: 140 NaCl, 4 KCl, 2 MgCl2, 4 glucose, 10 HEPES, 1 EGTA), cells were incubated with 200 µl Ca^2+^-free ECS for 10 minutes prior to imaging. A gravity-fed perfusion system was used to superfuse cells with ECS or test compounds at approximately 0.5 ml/min during imaging. The glass-bottom dishes were then mounted on an inverted Nikon Eclipse TE-2000S microscope and visualised with brightfield illumination (10x magnification).

Following excitation of Fluo-4-AM with a 470 nm LED (Cairn Research), intracellular Ca^2+^ mobilisation was captured on camera (Retiga Electro, Photometrics, AZ, USA) at 2.5 frames per second and 100 ms exposure time. Fluorescence was recorded at 520 nm using Micro-manager software (v1.4; NIH). ECS was washed over cells for 10 seconds prior to drug application (baseline), and for 4 minutes between drug applications (recovery). At the end of each protocol, 50 mM KCl was applied for 10 seconds to identify neurons and allow for normalisation of fluorescence.

### Ca^2+^ imaging data analysis

Fiji/ImageJ (NIH, MA, USA) was used to circle individual neurons and compute fluorescence intensity values (F) for each region of interest. Subsequently, a custom R script (RStudio, MA, USA) was used to identify responders and non-responders. From this, baseline-corrected normalised intensity values (F/F_pos_) could be generated based on the maximal fluorescence elicited by 50 mM KCl stimulation (F_pos_). Neurons were deemed responders to a drug challenge if a fluorescence increase of ≥0.1 F/F_pos_ was detected during drug application. For each experiment, the proportion of responding neurons and magnitude of the response was compared across control and treated groups.

### Magnetic-activated cell sorting of cultured sensory neurons

Magnetic-activated cell sorting (MACS) was used to generate pure DRG neuronal cultures that lacked non-neuronal cells, such as glia, thus enabling direct effects of IL-17 on DRG neurons to be determined. MACS equipment was purchased from Miltenyi Biotec and protocol developed according to previous studies (Higham et al., 2024; Thakur et al., 2014).

Thoracolumbar DRG from three mice were collected and cultured as described above, with the exception that the trypsin incubation step was omitted. Instead, DRG were incubated with collagenase (1 mg/ml with 6 mg/ml BSA) for 45 minutes at 37°C. Dulbecco’s phosphate-buffered saline (DPBS) was used to wash the cells before adding 120 µl MACS rinsing solution containing 0.5% (w/v) BSA along with 30 µl biotin-conjugated non-neuronal antibody cocktail for 5 minutes at 4°C. Samples were spun down (0.5 *g*, 7 min) and supernatant removed. Solutions were then incubated for 10 minutes at 4°C with 120 µl MACS rinsing solution containing 0.5% (w/v) BSA and 30 µl biotin-binding magnetic beads. The cell suspension was topped up to 500 µl with rinsing solution and gravity-fed through a magnetic column primed with 2.5 ml MACS buffer solution. Repeat additions of 1 ml rinsing solution was passed through the column to collect the 5 ml pure-DRG elute. This negative fraction was spun down (0.5 *g*, 7 min) and resuspended in Leibovitz L-15 medium (supplemented with GlutaMAX supplement, 2.6% (v/v) NaHCO_3_, 10% (v/v) FBS, 1.5% (v/v) glucose and P/S) prior to plating on laminin coated 35 mm poly-D-lysine-coated glass bottom dishes and incubated at 37°C for 3 hours to enable cell adhesion. Subsequently, the dishes were flooded with dissociation media and incubated for 48 hours (37°C, 5% CO_2_). Media was changed after 24 hours.

### Immunocytochemistry of cultured DRG neurons

DRG sensory neurons were cultured as above and plated onto laminin and poly-L-lysine coated coverslips. Following 48-hour incubation, cells were fixed in 4% PFA for 10 minutes and washed in phosphate-buffered saline (PBS). Cell were permeabilised with Triton-X (0.05%) for 5 minutes at room temperature (RT) prior to washing with PBS and incubation with blocking solution (1% goat serum in 0.2% Triton-X100) for 30 minutes. Subsequently, cells were incubated with a rabbit anti-βIII-tubulin primary antibody (1:500, Abcam: ab18207; RRID: AB_444319) overnight at 4°C. Cells were then washed in PBS and incubated with an Alexa Fluor-568 goat anti-rabbit secondary antibody (1:1000, Invitrogen: A11008; RRID:AB_143165) and 4’-6-diamidino-2-phenylindole (DAPI; 1:1000, Abcam) (1 hour, RT). Following a final wash in PBS (10 minutes) coverslips were mounted cell side down on glass slides using Mowiol 4-88 mounting medium (Sigma-Aldrich: 81381) and refrigerated for 30 minutes before imaging.

An Olympus BX51 microscope was used to image the slides. Fluorophores, namely DAPI and AlexaFluor-568, were excited with light sources at 350 nm and 568 nm, respectively. Images were then obtained using a Qicam CCD camera (QImaging) with a 100 ms exposure time and false coloured (DAPI, blue; βIII-tubulin, green). Omission of the primary antibody results in no βIII-tubulin staining (data not shown).

### Image analysis

Images were analysed using ImageJ as previously described (Hartig, 2013). An automatic ‘minimum error’ threshold algorithm was employed on 8-bit images of βIII-tubulin or DAPI staining to differentiate between background and regions of interest. Then binary and raw images underwent manual comparison, and the threshold adjusted manually to encompass all regions of interest. The threshold was placed at the first minimum after the major peak of the image histogram. Subsequently, touching objects in binary images were separated using the ImageJ watershed method. The identified objects, positive for either βIII-tubulin or DAPI, were automatically counted and a ratio of βIII-tubulin-positive cells (neurons) to DAPI-positive cells (neurons and non-neuronal cells) calculated.

### *Ex Vivo* electrophysiology lumbar splanchnic nerve recordings

Mice were euthanised as described above and the distal colon and associated nerve fibres (lumbar splanchnic nerve, LSN) isolated via laparotomy and transferred to a recording chamber (maintained at 32-35°C). The colorectum was cannulated and perfused luminally and serosally, at 100 µl/ml and 7 ml/min, respectively, with carbogenated Krebs solution (in mM: 124 NaCl, 4.8 KCl, 1.3 NaH_2_PO_4_, 25 NaHCO_3_, 1.2 MgSO_4_, 11.1 D-(+)-glucose and 2.5 CaCl_2_). To prevent smooth muscle contraction interfering with electrophysiological recordings, Krebs solution was supplemented with the muscarinic acetylcholine receptor antagonist, atropine, and voltage gated Ca^2+^ channel inhibitor, nifedipine (both at 10 µM).

Following cannulation, the LSN was carefully isolated and dissected free from its connective tissue and surrounding fat. Using a borosilicate glass suction electrode, the electrophysiological activity of the LSN was recorded in response to ramp distensions and drug challenges, as detailed in the provided protocols. Recordings were amplified, bandpass filtered (gain 5K; 100-1300 Hz; Neurolog, Digitimer Ltd, UK) and digitized at 20kHz (micro1401, CED, UK). Spike2 software was used for signal visualisation after digital filtering for 50 Hz noise (Humbug, Quest Scientific, Canada).

### Electrophysiology Protocols

Preparations were left for ∼30 minutes to stabilise prior to repeat ramp distensions (0-80 mmHg). A pressure transducer (Neurolog model NL108) was used to monitor luminal pressure up to 80 mmHg, which is above the threshold for all known visceral mechanoreceptors (Hughes et al., 2009; Ness & Gebhart, 1988). Five ramp distensions were performed in total, at 15-minute intervals. Recombinant mouse IL-17 (50 ng/ml) was perfused intraluminally (Harvard Apparatus, Pump 11 Elite) between ramps 3 and 5 to test for cytokine-induced mechanical hypersensitivity. For experiments in the presence of A425619, SB203580 or wortmannin, preparations were pre-treated with the antagonist and perfusion continued throughout the test protocol.

### Electrophysiological Data Analysis

Lumbar splanchnic nerve firing was determined by calculating the number of spikes passing the threshold (set at twice the level of background noise) and binned to determine the average firing frequency every 10 seconds. Baseline LSN firing was subtracted from increases in nerve activity in response to ramp distension. Changes in neuronal activity were measured at 5 mmHg intervals; from this the area under the curve (AUC) and peak changes in nerve firing could be determined and compared across vehicle and IL-17 treated groups. The AUC and peak nerve firing of ramp 5 (post-drug application) is presented as a percentage of ramp 3 (pre-drug application) at 80 mmHg.

### Statistical analysis

A Shapiro-Wilk test was used to assess the normality of all datasets and appropriate statistical tests were then employed. A significance level of P ≤ 0.05 was selected. Mean ± standard error of the mean is presented for all data. For Ca^2+^ imaging analysis, ‘n’ denotes the total number of dishes, and ‘N’ indicates the overall number of animals from which these dishes originated. For electrophysiology analysis, ‘N’ indicates the number of animals. Statistical analyses were performed using GraphPad Prism (version 10.4.2, GraphPad Software, San Diego, CA, USA).

## Results

### *In silico* analysis reveals increased *IL17A* expression in colonic biopsies from people with ulcerative colitis and marked co-expression of *Il17ra* with *Trpv1* in colon-projecting sensory neurons

*In silico* analysis of transcript expression in colonic biopsies showed a 10-fold increase in *IL17A* mRNA in samples from people with ulcerative colitis (UC), compared to noninflamed controls (NI) (P = 4.6 × 10^−9^, one-way ANOVA with Benjamini-Hochberg post-hoc test; **Fig. 1A**; data redrawn from (Higham et al., 2024)). Other *IL17* subtypes did not show such upregulation in UC, nor in drug-naïve and drug-treated Crohn’s disease (CDN and CDT, respectively) and recurrent abdominal pain (RAP) patient biopsies. Furthermore, the IL-12 cytokine family member, IL-23, which plays an indispensable role in driving IL-17 expression (Buonocore et al., 2010; Langrish et al., 2005), also showed heightened expression in UC biopsies compared to NI controls (P = 3.0 × 10^−6^, one-way ANOVA with Benjamini-Hochberg post-hoc test; **Fig. 1B**). However, *Il23r* is not expressed in murine colonic sensory neurons (**Figs. 1C-D**; data redrawn from (Hockley et al., 2019)). IL-23 is therefore unlikely to directly contribute to pain processing in colitis but may instead influence it indirectly through IL-17.

**Figure 1.**
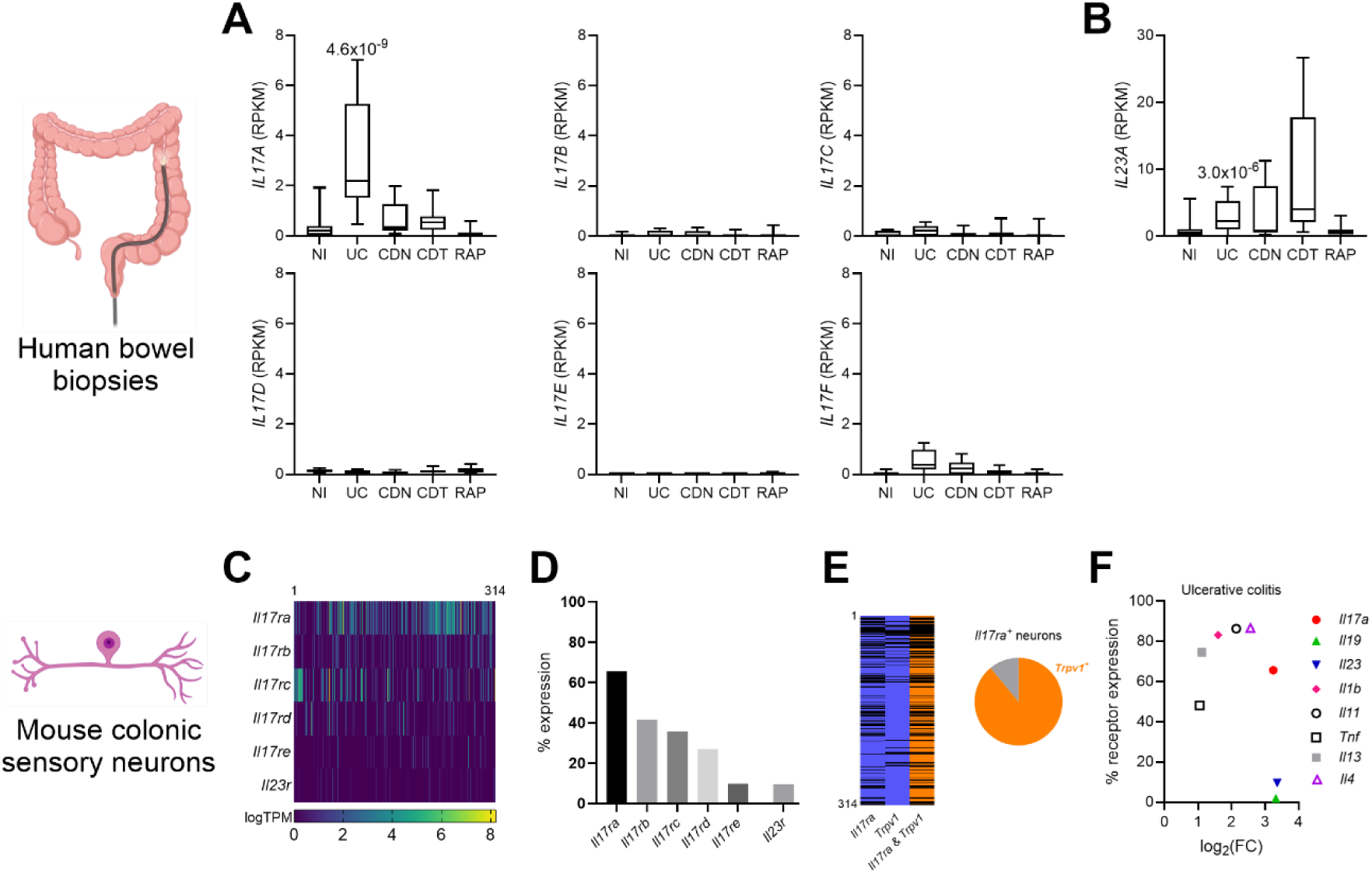
Upregulation of *IL17A* expression in the inflamed bowel and co-expression with *Trpv1* in colonic sensory neurons. (A) Expression (RPKM) of *IL17A*, *IL17B*, *IL17C*, *IL17D*, *IL17E* and *IL17F* in colonic biopsies taken from patients with no inflammation (NI), ulcerative colitis (UC), drug-naive Crohn’s disease (CDN), drug-treated Crohn’s disease (CDT) and recurrent abdominal pain (RAP). Median, interquartile range (box), and full range (whiskers) is shown (*IL17A* UC vs NI: P = 4.6 × 10^−9^, one-way ANOVA (main effect: P = 2.5 × 10^−9^; F(4, 58) = 17.14) with Benjamini-Hochberg post-hoc test (t = 6.89)). (B) Expression (RPKM) of *IL23A* in colonic biopsies taken from patients with NI, UC, CDN, CDT and RAP (*IL23A* UC vs NI: P = 3.0 × 10^−6^, one-way ANOVA (main effect: P = 1.1 × 10^−5^; F(4, 58) = 8.89) with Benjamini-Hochberg post-hoc test (t = 5.18)). Median, interquartile range (box), and full range (whiskers) is shown. Data redrawn from (Higham et al., 2024). (C) Heatmap showing the expression (logTPM) of *Il17ra*, *Il17rb*, *Il17rc*, *Il17rd, Il17re* and *Il23r* in murine colonic sensory neurons (data redrawn from Hockley et al., 2019). (D) Proportion of murine colonic sensory neurons expressing *Il17ra*, *Il17rb*, *Il17rc*, *Il17rd, Il17re* and *Il23r* (data redrawn from Hockley et al., 2019). (E) (Left) Heatmap showing the expression of *Il17ra* and *Trpv1* in murine colonic sensory neurons (data redrawn from Hockley et al., 2019). Neurons expressing either *Il17ra* or *Trpv1* are shown in blue, and neurons co-expressing *Il17ra* and *Trpv1* are shown in orange. (Right) Chart showing the fraction of *Il17ra*-expressing neurons which co-express *Trpv1* (orange). (F) Expression of selected cytokines in UC (given as log_2_FC relative to the NI group) against the fraction of murine colonic sensory neurons which express a putative receptor for each cytokine.

Among the five IL-17 receptor subtypes, *Il17ra* demonstrates the highest expression in murine colonic sensory neurons (**Fig. 1C-D**), with 65.6% of neurons expressing this receptor. Consistent with IL-17A binding a heterodimer of IL-17RA and IL-17RC (Toy et al., 2006), *Il17rc* also demonstrates notable expression in colonic sensory neurons (35.7%). Potential pronociceptive mediators in UC were identified by examining biopsy gene expression and cognate receptor expression in (murine) sensory neurons. The elevated IL-17 levels in UC biopsies, as well as the proportion of colonic sensory neurons expressing *Il17ra,* surpasses that of several other well-characterised proinflammatory cytokines (**Fig. 1F**). Additionally, the substantial proportion of *Il17ra*-positive sensory neurons co-expressing *Trpv1* (**Fig. 1E**), further supports a pronociceptive role for IL-17 in colitis.

### IL-17 increases [Ca^2+^]_i_ in capsaicin-sensitive DRG sensory neurons

Given the expression of *Il17ra* and *Il17rc* in mouse colonic sensory neurons, we sought to investigate the effects of IL-17 on intracellular Ca^2+^ mobilisation ([Ca^2+^]_i_) in cultured thoracolumbar DRG neurons. IL-17 increased [Ca^2+^]_i_ (**Figs. 2A-B**) in DRG neurons in a concentration-dependent manner with the proportion of neurons responding to IL-17 increasing from 7.94 ± 1.87% at 1 ng/ml to 15.80 ± 5.69% at 300 ng/ml (**Fig. 2C**), and responses to 100 and 300 ng/ml being significantly greater than to vehicle (vehicle vs 100 ng/ml IL-17: P = 0.0054; vehicle vs 300 ng/ml IL-17: P = 0.0152, Kruskal-Wallis test with Dunn’s multiple comparison; n = 5-8, N = 4-8; **Figs. 2C-D**). No significant differences in the magnitude of response in responding neurons were observed across the different IL-17 concentrations compared with the vehicle group (vehicle vs 300 ng/ml IL-17: P = 0.2750, ordinary one-way ANOVA; n = 5-8, N = 4-8, **Fig. 2E**). Furthermore, the soma area of IL-17-sensitive neurons (n = 48 cells) was significantly smaller than non-responders (n = 351 cells) (P = 0.0133, Mann-Whitney test. **Fig. 2F**), and consistent with IL-17 preferentially activating small neurons, 48.96% of IL-17 responders were co-sensitive to capsaicin (300 ng/ml group, **Figs. 2H-J**), suggesting that IL-17 can activate both capsaicin-sensitive and -insensitive neurons.

**Figure 2.**
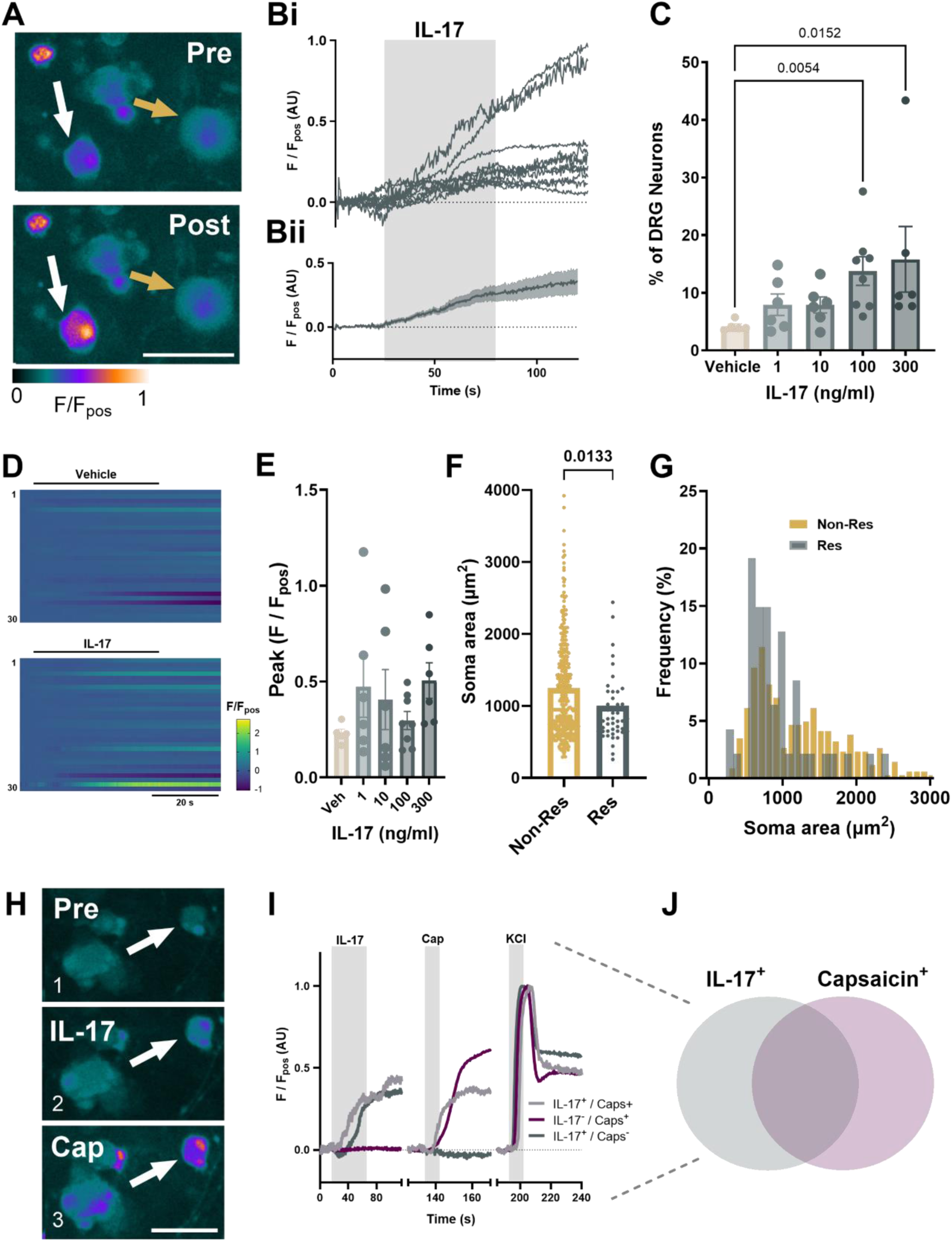
IL-17-evoked [Ca^2+^]_i_ increase in DRG neurons. (A) False-coloured images and corresponding individual (Bi) and averaged (Bii) traces showing Fluo4 fluorescence pre-and post-300 ng/ml IL-17 application (ten randomly selected IL-17-sensitive neurons used in Bi traces). White and yellow arrows highlight exemplar IL-17-sensitive and IL-17-insensitive neurons, respectively. Scale bar: 50 µm. (C) The proportion of DRG neurons responding to vehicle and 1 minute 1-300 ng/ml IL-17 stimulation (vehicle vs 100 ng/ml IL-17: P = 0.0054; vehicle vs 300 ng/ml IL-17: P = 0.0152, Kruskal-Wallis test (main effect: P = 0.0108; H = 13.09) with Dunn’s multiple comparison (Z = 3.204 and 2.895, respectively), n = 5-8, N = 4-8). (D) Heatmap showing Fluo4 fluorescence during the application of vehicle (top) or IL-17 (bottom) for 30 randomly selected neurons for 1 minute. (E) The averaged peak response of neurons responding to IL-17 (P = 0.3780, ordinary one-way ANOVA (F(4, 26) = 1.099), with Dunnett’s multiple comparisons test (n = 5-8, N = 4-8). (F) Scatter dot plot of average soma areas (µm^2^) (P = 0.0133, two-tailed Mann-Whitney test; U = 6574) and (G) cell size frequency distribution of responders and non-responders to IL-17. (H) False-coloured images of a neuron pre-(1) and post-IL-17 (2) and capsaicin (3) application. Scale bar: 50 µm. (I) Representative traces of capsaicin-sensitive and insensitive IL-17 responders and (J) Venn diagram depicting the co-sensitivity of neurons to IL-17 and capsaicin (300 ng/ml group, n = 6, N = 6).

Primary DRG neuron cultures contain non-neuronal cells, including glia, fibroblasts and immune cells (Bhuiyan et al., 2024). To determine if IL-17 directly stimulates sensory neurons, pure DRG neuronal cultures were generated using MACS (P < 0.0001, unpaired t test, n = 4-5 dishes, N = 3-4 pooled cultures, **Figs. 3A-C**), which showed a significant decrease in cell size (P < 0.0001, Mann-Whitney test, **Figs. 3D-E**) compared to unsorted cultures, consistent with a loss of large diameter neurons as previously reported for MACS (Higham et al., 2024; Thakur et al., 2014). The response to IL-17 was still observed in a comparable proportion of neurons following the removal of non-neuronal cells (P = 0.0986, unpaired t test, **Fig. 3G**) and the magnitude of IL-17 response was unchanged (P = 0.4530, unpaired t test, **Fig. 3H**). This confirms that IL-17 can directly activate sensory neurons, consistent with reported expression of IL-17 receptors in DRG neurons (Hockley et al., 2019).

**Figure 3.**
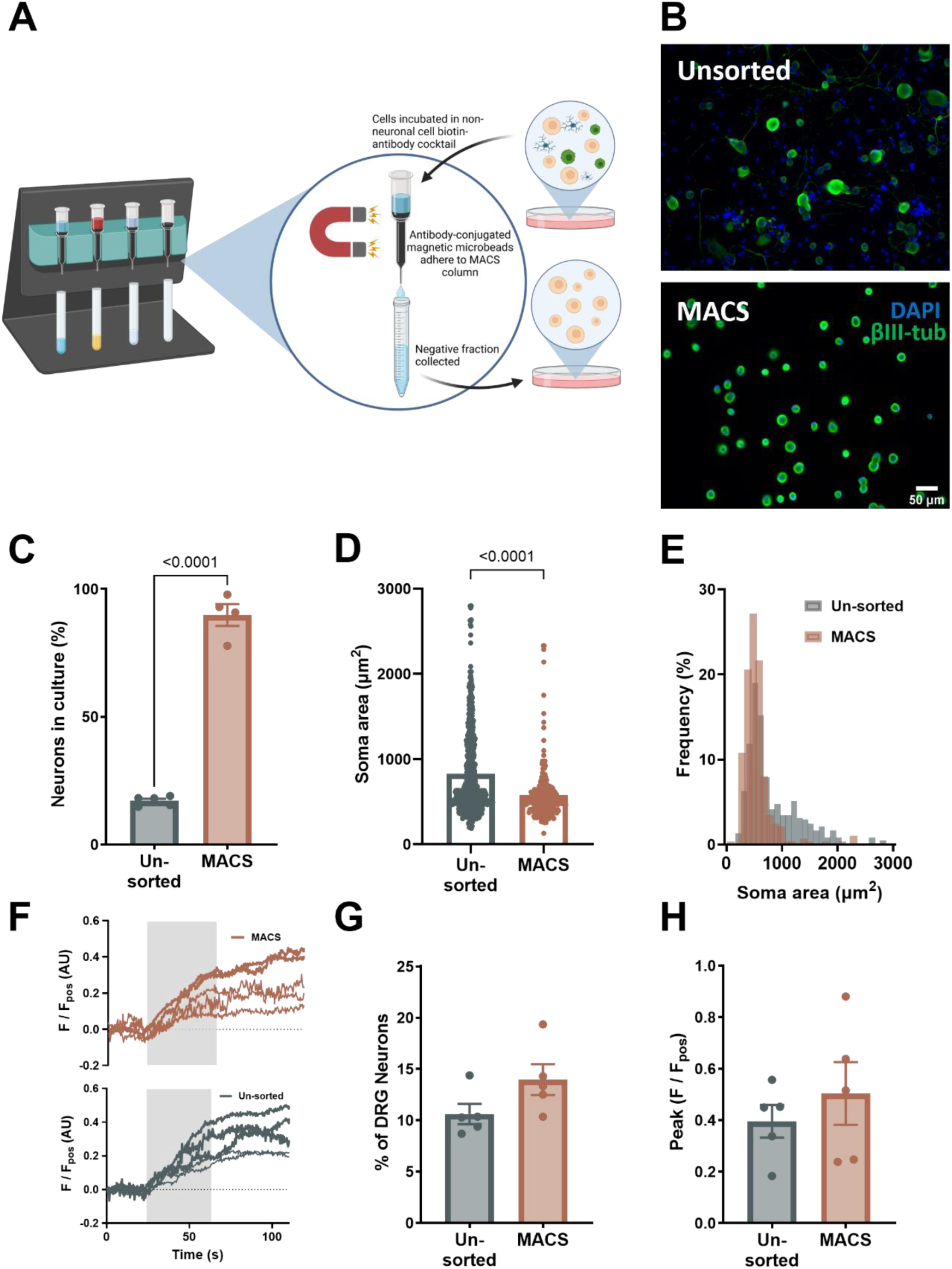
IL-17 directly stimulates DRG sensory neurons. (A) Schematic of MACS primary DRG neuron cultures. Cells are incubated in non-neuronal cell biotin-antibody cocktail. Antibody-conjugated magnetic microbeads adhere to the MACS column and the negative fraction collected is a pure DRG neuronal culture. (B) Example immunofluorescence images of nuclear DAPI stain (blue) and βIII-tubulin (green) in unsorted (*top*) and MACS (*bottom*) groups. (C) Proportion of neurons in culture in unsorted (n = 5 dishes from N = 3 pooled cultures) and MACS (n = 4 dishes from N = 4 pooled cultures) preparations (P < 0.0001, two-tailed unpaired t test; t, df = 18.78, 7). Enrichment of smaller-sized neurons demonstrated by (D) scatter dot plot of average soma areas (µm^2^) (P = <0.0001, two-tailed Mann-Whitney test; U = 60564) and (E) frequency of neurons of given soma areas for unsorted and MACS cultures. (F) Exemplar fluorescence trace showing the response to 100 ng/ml IL-17 in 5 randomly selected neurons in unsorted (bottom) and MACS (top) DRG cultures. (G) Proportion of IL-17-sensitive neurons in unsorted (n = 5 dishes from N = 3 pooled cultures) and MACS (n = 5 dishes from N = 4 pooled cultures) preparations (P = 0.0986, two-tailed unpaired t-test; t, df = 1.869, 8) and (H) peak amplitude of responders (P = 0.4530, two-tailed unpaired t-test; t, df = 0.7887, 8).

### IL-17-mediated Ca^2+^ signals in sensory neurons are dependent on extracellular Ca^2+^, TRPV1 signalling and activation of PI3K

To determine the source of Ca^2+^ contributing to the increase in [Ca^2+^]_i_ following IL-17 stimulation, responses were first examined in the absence of extracellular Ca^2+^ (**Figs. 4A-B**). In a Ca^2+^-free environment, the proportion of IL-17 responsive DRG neurons was significantly reduced (P = 0.0102, ordinary one-way ANOVA with Dunnett’s multiple comparison; n = 5, N = 5; **Fig. 4C**). Although not statistically significant, there was also a trend toward a reduced magnitude of IL-17-elicited responses (**Figs. 4D-E**). Consistent with this, depletion of intracellular Ca^2+^ stores by the sarco(endo)plasmic reticulum Ca^2+^-ATPase (SERCA) pump inhibitor, thapsigargin, did not significantly reduce the proportion of neurons responding to IL-17 (P = 0.3329, ordinary one-way ANOVA with Dunnett’s multiple comparison; n = 5, N = 5; **Fig. 4C**) or the magnitude of response (P = 0.9254, ordinary one-way ANOVA with Dunnett’s multiple comparison; n = 5, N = 5; **Fig. 4D**). These results suggest that IL-17-induced [Ca^2+^]_i_ mobilisation is driven by external Ca^2+^ entry.

**Figure 4.**
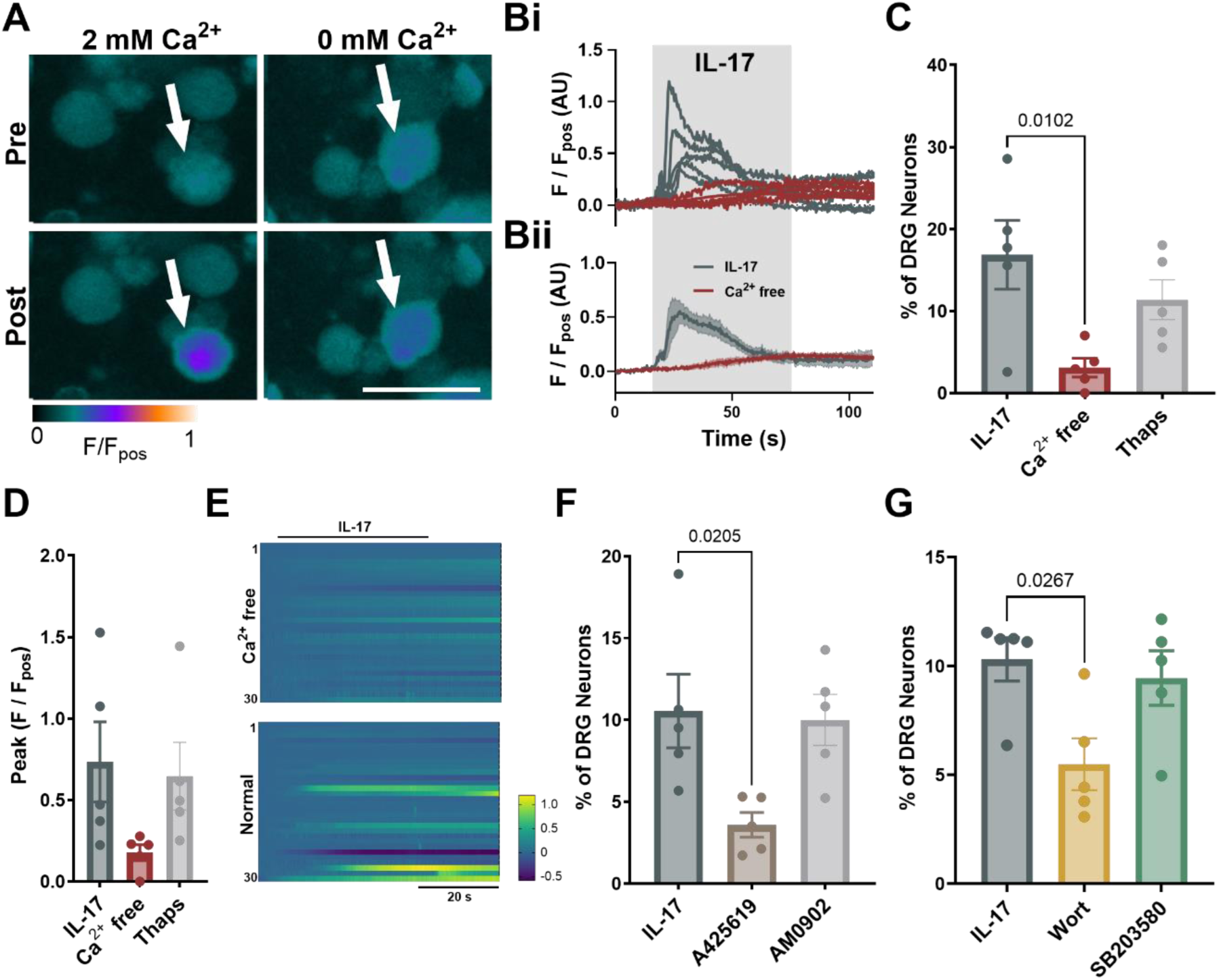
Activation of DRG sensory neurons by IL-17 is dependent on external Ca^2+^ entry, TRPV1 and PI3K. (A) False-coloured images and corresponding individual (Bi) and averaged (Bii) traces (5 randomly selected IL-17-sensitive neurons) showing Fluo-4 fluorescence Pre- and Post-IL-17 application in Ca^2+^-free (red) and normal (dark grey) ECS. White arrows highlight cells responding to IL-17. Scale bar: 50 µm. The proportion of DRG neurons responding (C) and peak response (D) to IL-17 following restriction of external Ca^2+^ (Proportion: P = 0.0102 (main effect: P = 0.0175; F(2, 12) = 5.779); Peak: P = 0.1023 (main effect: P = 0.1206; F(2, 12) = 2.537); ordinary one-way ANOVA with Dunnett’s multiple comparison; n = 5 from N = 5 independent cultures) or in the presence of thapsigargin (Thaps; SERCA pump inhibitor) (Proportion: P = 0.3329; Peak: P = 0.9254; ordinary one-way ANOVA with Dunnett’s multiple comparison; n = 5 from N = 5 independent cultures). (E) Heatmap showing Fluo-4 fluorescence during the application of IL-17 in Ca^2+^-free or normal ECS for 30 randomly selected neurons. (F) The proportion of DRG neurons responding to IL-17 following incubation with A425619 (TRPV1 inhibitor) or AM0902 (TRPA1 inhibitor) (P = 0.0205 and P = 0.9614, respectively, ordinary one-way ANOVA (main effect: P = 0.0196; F(2, 12) = 5.553) with Dunnett’s multiple comparison; n = 5, N = 5). (G) The proportion of DRG neurons responding to IL-17 following incubation with wortmannin (Wort; PI3K inhibitor) or SB203580 (P38 MAPK inhibitor) (P = 0.0267 and P = 0.9590, Kruskal-Wallis test (main effect: P = 0.0312; H = 6.5) with Dunn’s multiple comparison; n = 5, N = 4-5).

TRPV1 and TRPA1 are key Ca^2+^-permeable ion channels expressed by sensory neurons with established roles in sensory neuron activation that could be crucial to IL-17-induced nociception. High co-expression of IL-17RA and TRPV1 (**Fig. 1E**), along with previous studies identifying TRPV1 as a key player in IL-17 signalling (Luo et al., 2021), suggests that IL-17-induced Ca^2+^ entry may be TRPV1-mediated. In line with this, the TRPV1 inhibitor, A425619, significantly reduced the proportion of DRG neurons responding to IL-17 (P = 0.0205, ordinary one-way ANOVA with Dunnett’s multiple comparison; n = 5, N = 5; **Fig. 4F**). In contrast, no change in the proportion of neurons responding to IL-17 were observed following incubation with the TRPA1 inhibitor, AM0902 (P = 0.9614, ordinary one-way ANOVA with Dunnett’s multiple comparison; n = 5, N = 5; **Fig. 4F**). These results suggest that TRPV1, but not TRPA1, is essential for IL-17-induced activation of DRG sensory neurons.

To better understand the mechanism through which IL-17R signalling contributes to nociception in the GI tract, the role of key downstream signalling kinases, P38 MAPK and PI3K, was investigated. Interestingly, pre-incubation with the PI3K inhibitor wortmannin but not the P38 MAPK inhibitor SB203580, significantly reduced the percentage of DRG neurons responding to IL-17 (P = 0.0267 and P = 0.9590, Kruskal-Wallis test with Dunn’s multiple comparison; n = 5, N = 4-5; **Fig. 4G**), suggesting that PI3K may be recruited downstream of IL-17 receptor activation in DRG neurons.

### IL-17 sensitises colonic afferent responses to noxious distension via PI3K

Having established that IL-17 directly activates DRG sensory neurons, we next sought to determine if this effect translated to the activation and/or sensitisation of colonic afferents to noxious colorectal distension using *ex vivo* electrophysiology of LSN activity.

Stable responses to colonic distension (0 to 80 mmHg) were achieved following 2-3 distensions. For this reason, Ramp 3 was used as a control ramp in each protocol, after which tissues were perfused with 50 ng/ml IL-17 and the effects on the LSN response to mechanical distension investigated (**Fig. 5A**). Results are presented as the response to Ramp 5 as a % of Ramp 3 (at 80 mmHg). While IL-17 did not alter basal LSN activity (P = 0.3486, paired t test, N = 7-8; data not shown), incubation with IL-17 caused a marked increase in the afferent response to distension at noxious distending pressures (>20 mmHg) (P = 0.0094, Two-way repeated measures ANOVA, N = 7-8; **Fig. 5B**). Consistent with this, the difference in AUC, a readout of total afferent activity during colonic distension, of Ramp 5 normalised to Ramp 3 was significantly higher in IL-17 treated preparations compared to vehicle (P = 0.0037, Mann-Whitney test, N = 7-8; **Fig. 5C**), and the peak afferent firing in response to noxious ramp distension was elevated following treatment with IL-17 compared with vehicle (P = 0.0079, unpaired t test, N = 7-8; **Fig. 5D**).

**Figure 5.**
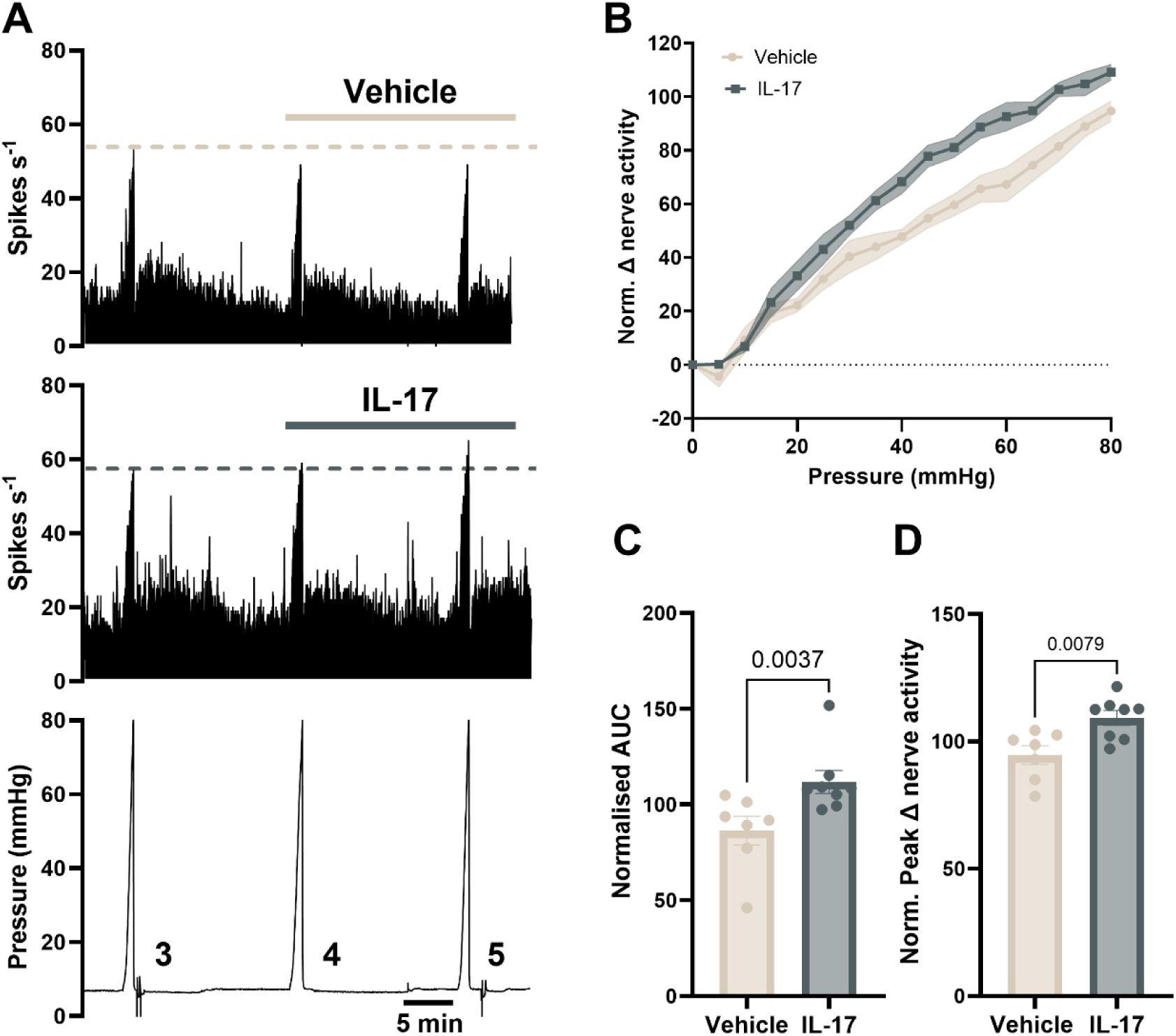
IL-17 evokes colonic afferent hypersensitivity to ramp distension. (A) Rate histograms and associated pressure trace of LSN recordings exemplifying afferent responses to ramp distensions 3-5 for vehicle and IL-17 (50 ng/ml) treated groups. (B) Afferent responses to ramp distension 5 (0-80 mmHg) in vehicle and IL-17 treated groups (normalised (norm.) to ramp distension 3) (main effect: P = 0.0094, vehicle vs. IL-17, two-way repeated measures ANOVA; F(1, 13) = 9.283; N = 7-8 per group). Shaded area is SEM. (C) Area under the curve (AUC) and (D) peak change in nerve activity (at 80 mmHg) of Ramp 5 (post-drug application) expressed as a percentage of Ramp 3 (pre-drug application) for vehicle and IL-17 treated preparations. Vehicle vs IL-17 compared with two-tailed Mann-Whitney test (P = 0.0037; U = 4) and two-tailed unpaired t-test (P = 0.0079; t, df = 3.136, 13), for AUC and peak change, respectively. All data represented as mean ± SEM.

Experiments were repeated in the presence of TRPV1, P38 MAPK or PI3K inhibitors (**Fig. 6**). Consistent with wortmannin’s ability to inhibit IL-17-mediated increases in [Ca^2+^]_i_ in sensory neurons (**Fig. 4G**), the PI3K inhibitor prevented IL-17-induced sensitisation of the colonic afferent response to noxious distension (Mean ± SEM IL-17: 117.1 ± 7.41; IL-17 + wortmannin: 92.33 ± 7.20; AUC: P = 0.0304; Peak: P = 0.0413, ordinary one-way ANOVA with Sidak’s multiple comparison; N = 8-9; **Figs. 6A-B**).

**Figure 6.**
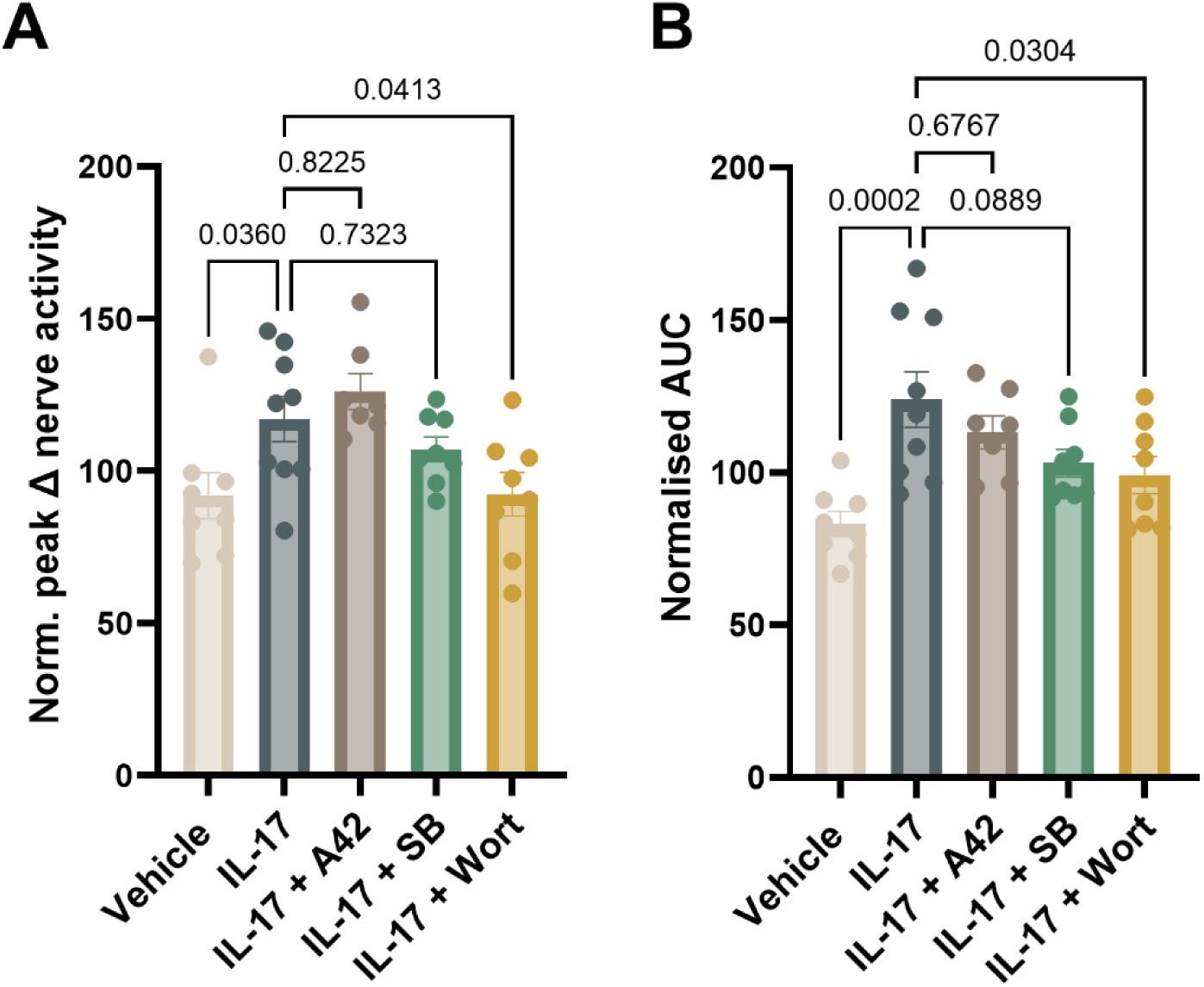
PI3K inhibition attenuates IL-17-induced mechanical hypersensitivity. (A) Peak nerve activity and (B) area under the curve (AUC) of Ramp 5 (post-drug application) expressed as a percentage of Ramp 3 (pre-drug application) for vehicle, IL-17 and antagonist-treated groups. Pharmacological inhibitors used: A42, A425619 (TRPV1 inhibitor); SB, SB203580 (P38 MAPK inhibitor); Wort, wortmannin (PI3K inhibitor). Comparisons were made to the IL-17 treated group using one-way ANOVA followed by Sidak’s multiple comparisons test (main effect: A: P = 0.0029; F(4, 35) = 4.946; B: P = 0.0008; F(4, 35) = 6.042) (N = 7-9 per group).

Pre-treatment with the TRPV1 antagonist had no significant effect on the response compared to IL-17 alone (AUC: P = 0.6767; Peak: P = 0.8225, ordinary one-way ANOVA with Sidak’s multiple comparison; N = 7-9; **Figs. 6A-B**). Similarly, no significant difference in IL-17-induced hypersensitivity was observed following incubation with the P38 MAPK inhibitor (AUC: P = 0.0889; Peak: P = 0.7323, ordinary one-way ANOVA with Sidak’s multiple comparison; N = 8-9; **Figs. 6A-B**), further corroborating our Ca^2+^ imaging data (**Fig. 4F**).

## Discussion

IL-17 is a proinflammatory cytokine associated with several inflammatory and autoimmune disorders, such as arthritis (Gaffen, 2009), psoriasis (Hawkes et al., 2018) and IBD (C. Abraham & J. Cho, 2009). Consistent with its well-documented role in IBD pathology, we observed an increase in IL-17A transcript levels in UC patients in line with previous reports (Fujino et al., 2003; Menesy et al., 2024), which, alongside the co-expression of its cognate IL-17 receptor subunits in colon projecting sensory neurons in mice, suggests that IL-17 potentially contributes to visceral nociceptor signalling in colitis. Here, we demonstrate a pronociceptive role for IL-17 in the colon, which was dependent on PI3K signalling, highlighting this cytokine and its downstream pathways as potential targets for treating pain associated with GI disease.

Findings from the current study confirmed that IL-17 evoked a rise in [Ca^2+^]_i_ in thoracolumbar DRG sensory neurons, in a proportion of neurons consistent with previous studies (Luo et al., 2021). These IL-17-sensitive neurons predominantly had smaller soma areas, and approximately half showed co-sensitivity to capsaicin, consistent with the activation of nociceptors and scRNA-seq data showing co-expression of *Trpv1* and *Il17ra* (Hockley et al., 2019). Although other cell types that reside within DRG, such as macrophages and glial cells (Bhuiyan et al., 2024), also express the receptor for IL-17 (Onishi & Gaffen, 2010), removal of non-neuronal cells with MACS confirmed that IL-17 was directly stimulating DRG sensory neurons. These findings are consistent with *Il17ra* and *Il17rc* expression in DRG neurons (Hockley et al., 2019) and add to growing evidence that IL-17 can directly signal to sensory neurons (Enamorado et al., 2023; Luo et al., 2019).

Building on this, we demonstrated that the IL-17-induced increase in [Ca^2+^]_i_ is mediated by external Ca^2+^ entry, most likely through TRPV1 given that responses were inhibited by pre-treatment with a TRPV1 inhibitor and Ca^2+^-free extracellular buffer. Furthermore, the response to IL-17 in DRG sensory neurons was attenuated following PI3K inhibition, consistent with previous reports of PI3K-mediated signalling in other cell types (Huang et al., 2007) and findings that PI3K and TRPV1 are functionally coupled (Bonnington & McNaughton, 2003; Stein et al., 2006). This work thus suggests that IL-17 may modulate TRPV1 signalling via its downstream modulation of PI3K in DRG neurons. Interestingly, unlike other proinflammatory cytokines such as TNF and IL-13, which have been shown to signal through P38 MAPK (Barker et al., 2023; Barker et al., 2022), pre-treatment with the P38 MAPK inhibitor SB203580 at comparable concentrations had no effect on the activation of DRG neurons by IL-17.

Given that IL-17 knockout mice demonstrate reduced mechanical hyperalgesia in several animal models of arthritis (Ebbinghaus et al., 2017; Richter et al., 2012) and IL-17 contributes to mechanical hypersensitivity following chemotherapy (Luo et al., 2019), we sought to translate our findings to the activation of colonic nociceptors by colorectal distension. Mechanical forces, such as stretching, are the relevant primary stimulus for pain generation in GI disease. Luminal perfusion of the colon with IL-17 induced a marked afferent mechanical hypersensitivity in responses to colorectal distension, which, consistent with our *in vitro* results in DRG neurons, was also blocked by PI3K inhibition. Interestingly, TRPV1 blockade did not reduce the mechanical sensitisation induced by IL-17, suggesting that IL-17-mediated mechanical hypersensitivity operates through TRPV1-independent mechanisms. This may reflect intrinsic differences between direct neuronal activation and sensitisation during mechanical distension, as TRPV1 is not primarily a mechanotransducer. Furthermore, in contrast to previous observations from our group that TNF and IL-13 sensitise colonic afferent mechanosensitivity via P38 MAPK signalling (Barker et al., 2023; Barker et al., 2022), pre-treatment with a P38 MAPK inhibitor had no effect on IL-17 mediated colonic afferent mechanosensitisation. Further work is needed to elucidate the molecular mechanisms by which IL-17 sensitises the colonic afferent response to noxious distension which appear to be distinct from those utilised by TNF and IL-13, with a key role for PI3K.

In summary, our study demonstrates that IL-17 directly activates DRG sensory neurons and contributes to mechanical sensitisation of colonic afferents. These effects are mediated through PI3K signalling, offering new insights into the role of IL-17 in pain processing. Our findings highlight a pro-nociceptive role for IL-17 signalling in the gut, emphasizing the potential of IL-17-targeted therapies for managing pain in GI diseases.

## Additional Information

### Author contributions

L.W.P. designed the research studies, conducted the experiments, acquired and analysed the data and wrote the manuscript. J.P.H. and K.H.B acquired and analysed the data. S.P. conceptualised the study and reviewed the manuscript. F.W., E.St.J.S., and D.C.B. designed the research studies and wrote the manuscript.

## Abbreviations

CD: Crohn’s disease
DRG: dorsal root ganglion
ECS: extracellular solution
GI: gastrointestinal
IBD: inflammatory bowel disease
IL-17: interleukin 17
IL-23: interleukin 23
LSN: lumbar splanchnic nerve
MACS: magnetic-activated cell sorting
PI3K: phosphoinositide 3-kinase
SERCA: sarco(endo)plasmic reticulum Ca^2+^-ATPase
TNF: tumour necrosis factor
TRPA1: transient receptor potential ankyrin 1
TRPV1: transient receptor potential vanilloid 1
UC: ulcerative colitis

## References

Abraham, C., & Cho, J. (2009). Interleukin-23/Th17 pathways and inflammatory bowel disease. Inflamm Bowel Dis, 15(7), 1090–1100. 10.1002/ibd.20894

Abraham, C., & Cho, J. H. (2009). Inflammatory bowel disease. N Engl J Med, 361(21), 2066–2078. 10.1056/NEJMra0804647

Barker, K. H., Higham, J. P., Pattison, L. A., Chessell, I. P., Welsh, F., Smith, E. S. J., & Bulmer, D. C. (2023). Sensitization of colonic nociceptors by IL-13 is dependent on JAK and p38 MAPK activity. Am J Physiol Gastrointest Liver Physiol, 324(4), G250–G261. 10.1152/ajpgi.00280.2022

Barker, K. H., Higham, J. P., Pattison, L. A., Taylor, T. S., Chessell, I. P., Welsh, F., Smith, E. S. J., & Bulmer, D. C. (2022). Sensitization of colonic nociceptors by TNFalpha is dependent on TNFR1 expression and p38 MAPK activity. J Physiol, 600(16), 3819–3836. 10.1113/JP283170

Bhuiyan, S. A., Xu, M., Yang, L., Semizoglou, E., Bhatia, P., Pantaleo, K. I., Tochitsky, I., Jain, A., Erdogan, B., Blair, S., Cat, V., Mwirigi, J. M., Sankaranarayanan, I., Tavares-Ferreira, D., Green, U., McIlvried, L. A., Copits, B. A., Bertels, Z., Del Rosario, J. S.,…Renthal, W. (2024). Harmonized cross-species cell atlases of trigeminal and dorsal root ganglia. Sci Adv, 10(25), eadj9173. 10.1126/sciadv.adj9173

Bonnington, J. K., & McNaughton, P. A. (2003). Signalling pathways involved in the sensitisation of mouse nociceptive neurones by nerve growth factor. J Physiol, 551(Pt 2), 433–446. 10.1113/jphysiol.2003.039990

Buonocore, S., Ahern, P. P., Uhlig, H. H., Ivanov, II, Littman, D. R., Maloy, K. J., & Powrie, F. (2010). Innate lymphoid cells drive interleukin-23-dependent innate intestinal pathology. Nature, 464(7293), 1371–1375. 10.1038/nature08949

Chang, S. H., & Dong, C. (2007). A novel heterodimeric cytokine consisting of IL-17 and IL-17F regulates inflammatory responses. Cell Res, 17(5), 435–440. 10.1038/cr.2007.35

Chang, S. H., Park, H., & Dong, C. (2006). Act1 adaptor protein is an immediate and essential signaling component of interleukin-17 receptor. J Biol Chem, 281(47), 35603–35607. 10.1074/jbc.C600256200

Ebbinghaus, M., Natura, G., Segond von Banchet, G., Hensellek, S., Bottcher, M., Hoffmann, B., Salah, F. S., Gajda, M., Kamradt, T., & Schaible, H. G. (2017). Interleukin-17A is involved in mechanical hyperalgesia but not in the severity of murine antigen-induced arthritis. Sci Rep, 7(1), 10334. 10.1038/s41598-017-10509-5

Enamorado, M., Kulalert, W., Han, S. J., Rao, I., Delaleu, J., Link, V. M., Yong, D., Smelkinson, M., Gil, L., Nakajima, S., Linehan, J. L., Bouladoux, N., Wlaschin, J., Kabat, J., Kamenyeva, O., Deng, L., Gribonika, I., Chesler, A. T., Chiu, I. M.,…Belkaid, Y. (2023). Immunity to the microbiota promotes sensory neuron regeneration. Cell, 186(3), 607–620 e617. 10.1016/j.cell.2022.12.037

Friedrich, M., Pohin, M., & Powrie, F. (2019). Cytokine Networks in the Pathophysiology of Inflammatory Bowel Disease. Immunity, 50(4), 992–1006. 10.1016/j.immuni.2019.03.017

Fujino, S., Andoh, A., Bamba, S., Ogawa, A., Hata, K., Araki, Y., Bamba, T., & Fujiyama, Y. (2003). Increased expression of interleukin 17 in inflammatory bowel disease. Gut, 52(1), 65–70. 10.1136/gut.52.1.65

Gaffen, S. L. (2009). The role of interleukin-17 in the pathogenesis of rheumatoid arthritis. Curr Rheumatol Rep, 11(5), 365–370. 10.1007/s11926-009-0052-y

Gaffen, S. L., Jain, R., Garg, A. V., & Cua, D. J. (2014). The IL-23-IL-17 immune axis: from mechanisms to therapeutic testing. Nat Rev Immunol, 14(9), 585–600. 10.1038/nri3707

Grundy, L., Erickson, A., & Brierley, S. M. (2019). Visceral Pain. Annu Rev Physiol, 81, 261–284. 10.1146/annurev-physiol-020518-114525

Hartig, S. M. (2013). Basic image analysis and manipulation in ImageJ. Curr Protoc Mol Biol, Chapter 14, Unit14 15. 10.1002/0471142727.mb1415s102

Hawkes, J. E., Yan, B. Y., Chan, T. C., & Krueger, J. G. (2018). Discovery of the IL-23/IL-17 Signaling Pathway and the Treatment of Psoriasis. J Immunol, 201(6), 1605–1613. 10.4049/jimmunol.1800013

Higham, J. P., Bhebhe, C. N., Gupta, R. A., Tranter, M. M., Barakat, F. M., Dogra, H., Bab, N., Wozniak, E., Barker, K. H., Wilson, C. H., Mein, C. A., Raine, T., Cox, J. J., Wood, J. N., Croft, N. M., Wright, P. D., & Bulmer, D. C. (2024). Transcriptomic profiling reveals a pronociceptive role for angiotensin II in inflammatory bowel disease. Pain, 165(7), 1592–1604. 10.1097/j.pain.0000000000003159

Hockley, J. R. F., Taylor, T. S., Callejo, G., Wilbrey, A. L., Gutteridge, A., Bach, K., Winchester, W. J., Bulmer, D. C., McMurray, G., & Smith, E. S. J. (2019). Single-cell RNAseq reveals seven classes of colonic sensory neuron. Gut, 68(4), 633–644. 10.1136/gutjnl-2017-315631

Huang, F., Kao, C. Y., Wachi, S., Thai, P., Ryu, J., & Wu, R. (2007). Requirement for both JAK-mediated PI3K signaling and ACT1/TRAF6/TAK1-dependent NF-kappaB activation by IL-17A in enhancing cytokine expression in human airway epithelial cells. J Immunol, 179(10), 6504–6513. 10.4049/jimmunol.179.10.6504

Hughes, P. A., Brierley, S. M., Martin, C. M., Brookes, S. J., Linden, D. R., & Blackshaw, L. A. (2009). Post-inflammatory colonic afferent sensitisation: different subtypes, different pathways and different time courses. Gut, 58(10), 1333–1341. 10.1136/gut.2008.170811

Langrish, C. L., Chen, Y., Blumenschein, W. M., Mattson, J., Basham, B., Sedgwick, J. D., McClanahan, T., Kastelein, R. A., & Cua, D. J. (2005). IL-23 drives a pathogenic T cell population that induces autoimmune inflammation. J Exp Med, 201(2), 233–240. 10.1084/jem.20041257

Lee, J. S., Tato, C. M., Joyce-Shaikh, B., Gulen, M. F., Cayatte, C., Chen, Y., Blumenschein, W. M., Judo, M., Ayanoglu, G., McClanahan, T. K., Li, X., & Cua, D. J. (2015). Interleukin-23-Independent IL-17 Production Regulates Intestinal Epithelial Permeability. Immunity, 43(4), 727–738. 10.1016/j.immuni.2015.09.003

Linden, A. (2007). A role for the cytoplasmic adaptor protein Act1 in mediating IL-17 signaling. Sci STKE, 2007(398), re4. 10.1126/stke.3982007re4

Luo, H., Liu, H. Z., Zhang, W. W., Matsuda, M., Lv, N., Chen, G., Xu, Z. Z., & Zhang, Y. Q. (2019). Interleukin-17 Regulates Neuron-Glial Communications, Synaptic Transmission, and Neuropathic Pain after Chemotherapy. Cell Rep, 29(8), 2384–2397 e2385. 10.1016/j.celrep.2019.10.085

Luo, X., Chen, O., Wang, Z., Bang, S., Ji, J., Lee, S. H., Huh, Y., Furutani, K., He, Q., Tao, X., Ko, M. C., Bortsov, A., Donnelly, C. R., Chen, Y., Nackley, A., Berta, T., & Ji, R. R. (2021). IL-23/IL-17A/TRPV1 axis produces mechanical pain via macrophage-sensory neuron crosstalk in female mice. Neuron, 109(17), 2691–2706 e2695. 10.1016/j.neuron.2021.06.015

McGeachy, M. J., Chen, Y., Tato, C. M., Laurence, A., Joyce-Shaikh, B., Blumenschein, W. M., McClanahan, T. K., O’Shea, J. J., & Cua, D. J. (2009). The interleukin 23 receptor is essential for the terminal differentiation of interleukin 17-producing effector T helper cells in vivo. Nat Immunol, 10(3), 314–324. 10.1038/ni.1698

Menesy, A., Hammad, M., Aref, S., & Abozeid, F. A. M. (2024). Level of interleukin 17 in inflammatory bowel disease and its relation with disease activity. BMC Gastroenterol, 24(1), 135. 10.1186/s12876-024-03218-7

Ness, T. J., & Gebhart, G. F. (1988). Colorectal distension as a noxious visceral stimulus: physiologic and pharmacologic characterization of pseudaffective reflexes in the rat. Brain Res, 450(1-2), 153–169. 10.1016/0006-8993(88)91555-7

Onishi, R. M., & Gaffen, S. L. (2010). Interleukin-17 and its target genes: mechanisms of interleukin-17 function in disease. Immunology, 129(3), 311–321. 10.1111/j.1365-2567.2009.03240.x

Pinto, L. G., Cunha, T. M., Vieira, S. M., Lemos, H. P., Verri, W. A., Jr., Cunha, F. Q., & Ferreira, S. H. (2010). IL-17 mediates articular hypernociception in antigen-induced arthritis in mice. Pain, 148(2), 247–256. 10.1016/j.pain.2009.11.006

Richter, F., Natura, G., Ebbinghaus, M., von Banchet, G. S., Hensellek, S., Konig, C., Brauer, R., & Schaible, H. G. (2012). Interleukin-17 sensitizes joint nociceptors to mechanical stimuli and contributes to arthritic pain through neuronal interleukin-17 receptors in rodents. Arthritis Rheum, 64(12), 4125–4134. 10.1002/art.37695

Robinson, D. R., McNaughton, P. A., Evans, M. L., & Hicks, G. A. (2004). Characterization of the primary spinal afferent innervation of the mouse colon using retrograde labelling. Neurogastroenterol Motil, 16(1), 113–124. 10.1046/j.1365-2982.2003.00456.x

Stein, A. T., Ufret-Vincenty, C. A., Hua, L., Santana, L. F., & Gordon, S. E. (2006). Phosphoinositide 3-kinase binds to TRPV1 and mediates NGF-stimulated TRPV1 trafficking to the plasma membrane. J Gen Physiol, 128(5), 509–522. 10.1085/jgp.200609576

Thakur, M., Crow, M., Richards, N., Davey, G. I., Levine, E., Kelleher, J. H., Agley, C. C., Denk, F., Harridge, S. D., & McMahon, S. B. (2014). Defining the nociceptor transcriptome. Front Mol Neurosci, 7, 87. 10.3389/fnmol.2014.00087

Toy, D., Kugler, D., Wolfson, M., Vanden Bos, T., Gurgel, J., Derry, J., Tocker, J., & Peschon, J. (2006). Cutting edge: interleukin 17 signals through a heteromeric receptor complex. J Immunol, 177(1), 36–39. 10.4049/jimmunol.177.1.36

